# Version-aware question answering over living oncology guidelines

**DOI:** 10.64898/2026.10.04.756483

**Authors:** Tampu Ravi Kumar, Anirudh Iyengar Kaniyar Narayana Iyengar, Muhammad Ali Khan, Kaneez Zahra Rubab Khakwani, Karan Kathuria, M Bassam Sonbol, Irbaz Bin Riaz, Vivek Gupta

## Abstract

Living clinical guidelines revise recommendations as evidence emerges, so an AI answer can be faithful to a real guideline and still be out of date. Here we examine this failure using ASCOBench, 288 unique questions in 96 three-turn conversations grounded in versioned American Society of Clinical Oncology (ASCO) breast and prostate cancer guidelines, with oncologist-reviewed reference answers. Oncologists corrected one in four version-sensitive answers drafted by a frontier model, against fewer than one in ten factual ones. Across the guideline corpus, 19 recommendations changed between versions, seven of them reversals, yet only 1 of 14 superseded documents states that it has been replaced. With a strong model, retrieval over several guideline versions made stale answers four times more frequent than answering without retrieval. SentryLine, a verification-first system, lowered incorrect answers to 4.2-5.2% from 9.7-22.9% for baselines across three models. Recognizing change before a guideline declares it remains an open problem.

## 1 Introduction

Recommendations in oncology change as trials report, and the volume of new evidence far exceeds what any clinician can read: more than 1.5 million articles enter PubMed each year [1], and keeping up with every applicable guideline would by one estimate take a primary care physician 27 hours a day [2]. ASCO’s answer is the living guideline, revised on a rolling basis rather than at fixed intervals. A revision can do more than refine earlier advice; it can reverse it. When ASCO updated its 2016 breast cancer biomarker guideline in 2019 to incorporate TAILORx, it replaced an unqualified permission to use the 21-gene recurrence score with age-specific score thresholds for offering chemotherapy [3, 4], and its 2022 update reversed several of the 2019 recommendations (Section 2.3).

Clinicians increasingly consult AI tools at the point of care. In a 2026 survey of nearly 1,700 US physicians, 81% reported using AI in their practice [5], and OpenEvidence, a literature-grounded medical answer engine, told NBC News in May 2026 that about 65% of US physicians use it [6]. ASCO deployed its own Gemini-based Guidelines Assistant [7], and patients use general-purpose chatbots too: about a third of US adults turned to one for health information in the past year [8]. These tools present citations, which signals that an answer is grounded. A citation, however, does not guarantee that the cited guideline version is still current. Independent evidence on how these tools perform is limited and conflicting. In a study comparing OpenEvidence and UpToDate Expert AI with GPT-5.2, Gemini 3.1 Pro and Claude Opus 4.6, the general-purpose models outperformed both clinical tools on 500 MedQA questions, 500 HealthBench items and 100 real physician queries; on MedQA, Gemini reached 97.4% against 89.6% for OpenEvidence and 88.4% for UpToDate Expert AI, and on real queries the clinical tools performed comparably to Google’s search AI Overview [9]. A later preprint built on 620 real point-of-care queries from the OpenEvidence platform reached the opposite conclusion: 149 specialty-matched physicians preferred OpenEvidence over Claude Opus 4.8, Gemini 3.1 Pro and GPT-5.5 on accuracy, clinical utility, source quality, verifiability and completeness, with win margins of 25-39 percentage points [10]. The two studies differ in query source, grading design and sponsor involvement, and neither asks whether an answer reflects the current version of a guideline.

We call this failure mode *version staleness*: an answer that is faithful to a real guideline but to a superseded version of it. It differs from hallucination in a way that matters clinically. A hallucinated claim has no source and can be caught by checking the reference [11]; a stale claim has a legitimate source and passes that check. Medical language models that store clinical knowledge in their weights [12–15] answer from what was current when they were trained, and language models degrade on facts that have changed since [16, 17]. Retrieval-augmented generation [18] grounds answers in external documents and has been widely evaluated in medicine [19, 20], but guideline-grounded systems in other specialties treat each guideline as a static snapshot [21–23]. Errors in oncology are compounded by low health literacy among patients [24], yet existing evaluations of oncology AI use exam-style multiple-choice questions [25]. To our knowledge, no benchmark or evaluation links questions to explicit guideline versions with expert-reviewed answers, and no study has measured how often models return the superseded recommendation when a guideline has changed.

Here we examine version staleness in question answering over the American Society of Clinical Oncology (ASCO) breast and prostate cancer guidelines (Fig. 1), building on ASCOBench and SentryLine, a benchmark and a verification-first question-answering system that we described previously [26]. In ASCOBench, every question and reference answer is tied to a specific guideline version and page, and oncologists reviewed every answer drafted by a frontier model. We first analyse that review as an evaluation of model-generated clinical content, asking which kinds of question the model most often got wrong. We then catalogue the recommendations in the guideline corpus that changed between versions, the evidence each update cites and the time between versions, and ask whether superseded documents say that they have been replaced. Finally, after removing duplicated conversations from the test set, we re-analyse the answers of SentryLine and of retrieval and prompting baselines with three language models, adding conversation-level confidence intervals, an analysis of stale answers on version-sensitive questions and an assessment of whether the system’s verification verdicts identify answers that need review. In this study, we show that version-sensitive answers were the ones oncologists most often had to correct, that the guideline corpus contains 19 changed recommendations of which seven are reversals while only one of 14 superseded documents announces its replacement, and that verification against the current version lowered incorrect and stale answers with every model tested. SentryLine and an earlier version of ASCOBench were introduced as a system demonstration [26]; here the benchmark and its annotation are described in full, all outputs are re-analysed on a unique questions from the test set, and the census of recommendation changes is new. We also consider what these results imply for the harder problem of recognizing that a recommendation should change before a guideline declares it.

**Fig. 1.**
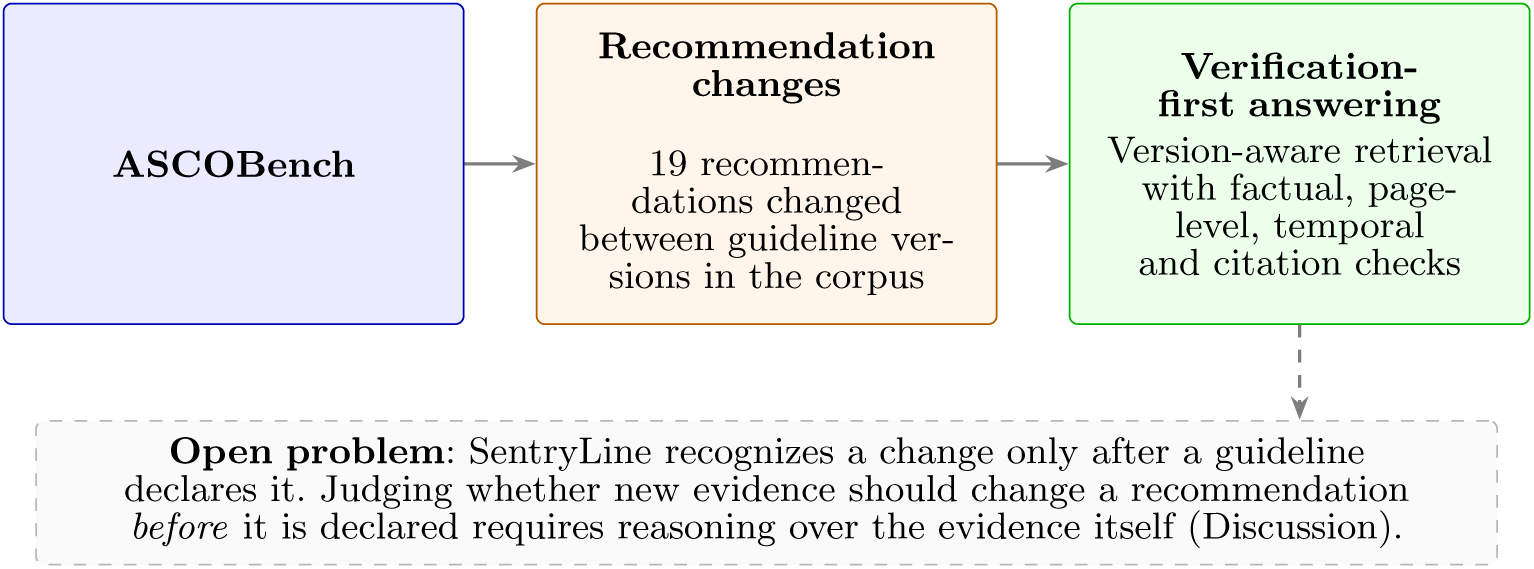
Overview of the study. An oncologist-reviewed benchmark tied to guideline versions measures how often answers rest on superseded guidance; a census of the guideline corpus shows how recommendations change and whether superseded documents say so; and SentryLine verifies answers against the current version once an update has been declared. The dashed box marks the problem this study does not address.

## 2 Results

### 2.1 A version-linked benchmark reviewed by oncologists

ASCOBench consists of three-turn conversations about breast and prostate cancer ASCO guidelines (Fig. 2). Each conversation probes one of four abilities that question answering over living guidelines requires: stating guideline facts precisely, reasoning across several recommendations, telling the current recommendation from a superseded one, and answering in terms the reader can use (Table 1). Questions and answers were drafted by a frontier model (Claude Opus 4.8) from the guideline text, each tied to a quotation on a specific page, and every answer in the test set was then reviewed by oncologists, who accepted, revised or rewrote it to produce the reference answer and scored the draft on seven clinical dimensions (Methods). After removing nine conversations that duplicated others, the test set used throughout this study holds 96 unique conversations and 288 turns (Table 2).

**Fig. 2.**
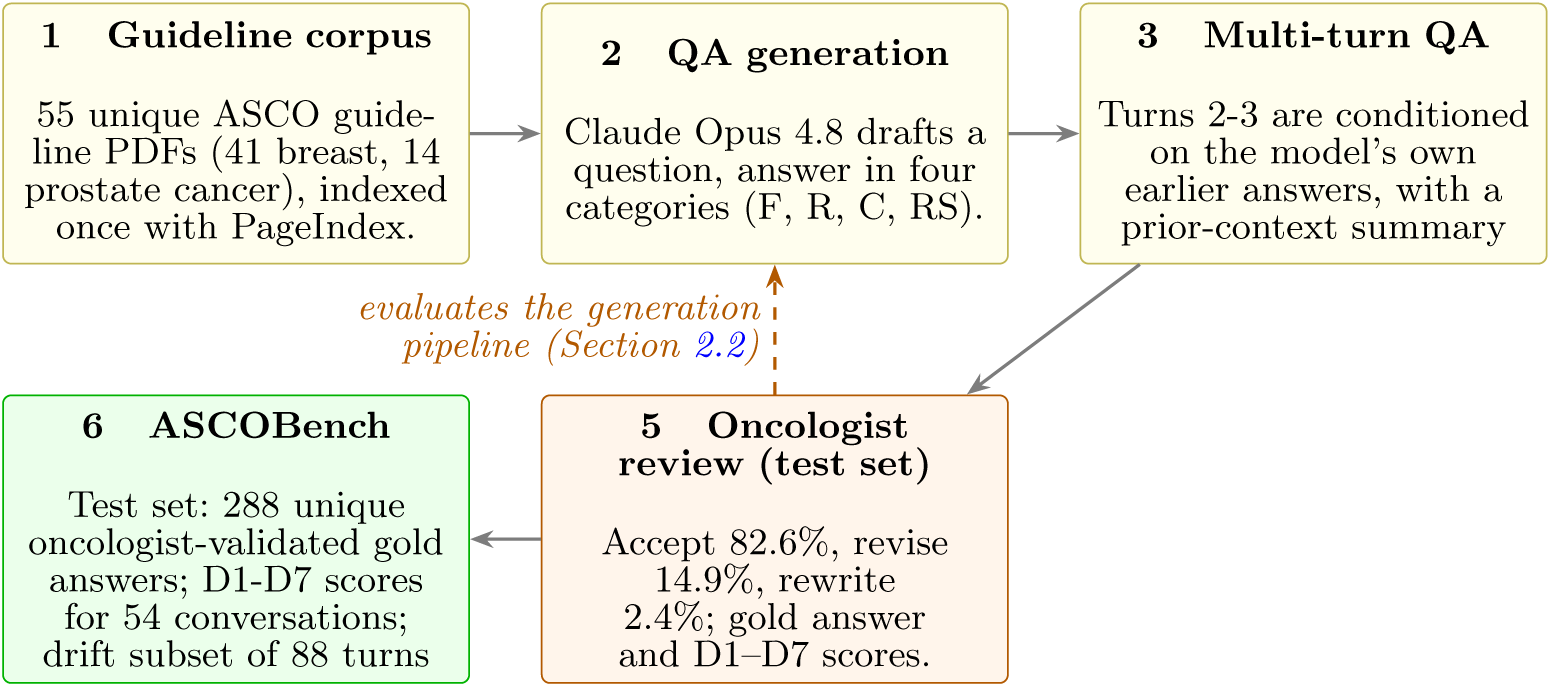
ASCOBench construction pipeline. Guideline documents are converted into grounded question-answer pairs (stages 1-2), expanded into three-turn conversations (3). Oncologists review every model-drafted answer in the test set (4), producing the gold references (5). Because every draft is reviewed, the review outcomes also measure how well the generation pipeline performs (dashed arrow). F, Factual; R, Reasoning; C, Contrasting; RS, Role-Specific.

**Table 1.** Question categories. Turn counts in the test split by target role. Target role selects the scoring anchors for role calibration (D4). Examples are turn-1 questions from the test set.

| Category | What it tests | Clinician /<br>patient turns | Example (turn 1) |
| --- | --- | --- | --- |
| Factual | Clinical thresholds, drug combinations, dosing and biomarker cutoffs stated in the guideline | 27 / 12 | “What is the dose of elacestrant used in the EMERALD trial?” |
| Reasoning | Step-by-step clinical logic that combines several recommendations rather than retrieving one fact | 54 / 39 | “Why is the Panel unable to recommend adjuvant denosumab for breast cancer recurrence prevention?” |
| Contrasting | An outdated standard named alongside the current recommendation, with a plausible decoy; tests retrieval of the current version | 60 / 48 | “Previously, the 2020 ASCO hereditary breast cancer guideline stated that radiation therapy should not be withheld in ATM carriers. Has this recommendation changed in subsequent updates?” |
| Role-Specific | Questions about what a recommendation means for a specific reader | 24 / 24 | “I have HER2-positive breast cancer that has spread to my brain. What are my options?” |

**Table 2.**
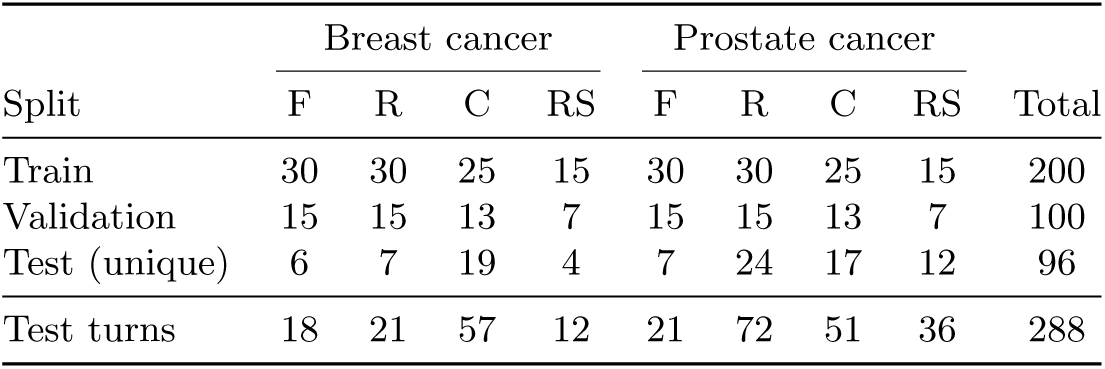
ASCOBench characteristics. Conversation counts by split, cancer type and question category (F, Factual; R, Reasoning; C, Contrasting; RS, Role-Specific). Test counts are unique conversations in the annotated test set, after removing nine exact duplicates; each conversation has three turns.

| Split | Breast cancer |  |  |  | Prostate cancer |  |  |  | Total |
| --- | --- | --- | --- | --- | --- | --- | --- | --- | --- |
|  | F | R | C | RS | F | R | C | RS |  |
| Train | 30 | 30 | 25 | 15 | 30 | 30 | 25 | 15 | 200 |
| Validation | 15 | 15 | 13 | 7 | 15 | 15 | 13 | 7 | 100 |
| Test (unique) | 6 | 7 | 19 | 4 | 7 | 24 | 17 | 12 | 96 |
| Test turns | 18 | 21 | 57 | 12 | 21 | 72 | 51 | 36 | 288 |

### 2.2 Oncologists corrected version-sensitive drafts most often

Because every drafted answer was reviewed, the review also measures how well the generation pipeline performs. Oncologists accepted 82.6% of the pipeline’s drafted answers as references without change, revised 14.9% and rewrote 2.4% (Table 3, Fig. 3). Errors concentrated where the benchmark is meant to be hard. Contrasting turns, which hinge on which guideline version is current, needed the most correction (19.4% revised, 5.6% rewritten, one in four overall), and Factual turns the least (92.3% accepted). The generator, in other words, struggled with the same version-tracking task that ASCOBench is designed to test. Patient-facing drafts were rarely rewritten (0.8%) but were revised more often than clinician-facing ones (17.9% vs 12.7%). Decisions and accuracy scores agreed in most rows but not all: 22 turns were revised yet scored fully correct, which we interpret as edits for completeness or style rather than corrections of substance, and 4 further turns had a decision inconsistent with their D1 score.

**Fig. 3.**
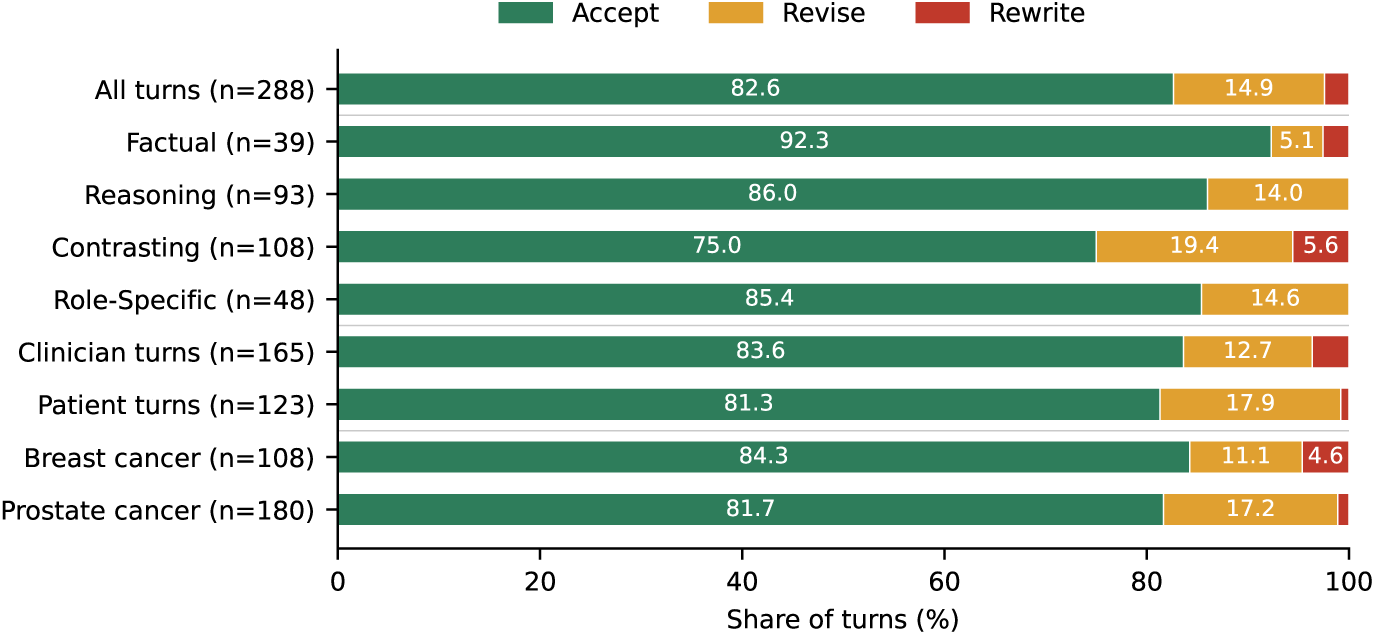
Reference-answer decisions by question category, target role and cancer type. Share of test turns whose model-drafted answer the oncologists accepted, revised or rewrote. Unlabelled segments are below 4%.

**Table 3.** Evaluation of the generation pipeline: oncologist decisions on drafted answers for all 288 test turns. Values are percentages of turns in each group.

| Group | Turns | Accept (%) | Revise (%) | Rewrite (%) |
| --- | --- | --- | --- | --- |
| All turns | 288 | 82.6 | 14.9 | 2.4 |
| Factual | 39 | 92.3 | 5.1 | 2.6 |
| Reasoning | 93 | 86.0 | 14.0 | 0.0 |
| Contrasting | 108 | 75.0 | 19.4 | 5.6 |
| Role-Specific | 48 | 85.4 | 14.6 | 0.0 |
| Clinician-facing | 165 | 83.6 | 12.7 | 3.6 |
| Patient-facing | 123 | 81.3 | 17.9 | 0.8 |
| Breast cancer | 108 | 84.3 | 11.1 | 4.6 |
| Prostate cancer | 180 | 81.7 | 17.2 | 1.1 |

The pipeline’s drafted answers were rated fully accurate in 86.2% of scored turns, partially correct in 11.3% and incorrect in 2.6% (D1; Table 4, Fig. 4). 8.2% were judged ungrounded in ASCO content (D2). None was judged unsafe (D3), and every scored follow-up turn retained the conversation’s context (D6). Role calibration (D4) was weaker for patient-facing turns, of which 11.1% were rated poorly calibrated and 72.2% fully calibrated, than for clinician-facing turns (2.2% and 84.4%). On the drift subset, 2 of 88 turns (2.3%) gave the superseded recommendation or conflated versions (D5). Of the 54 fully annotated conversations, 81.5% were rated fully coherent and one incoherent (D7). These scores describe the generation pipeline’s drafts, not the systems evaluated in Section 2.4.

**Fig. 4.**
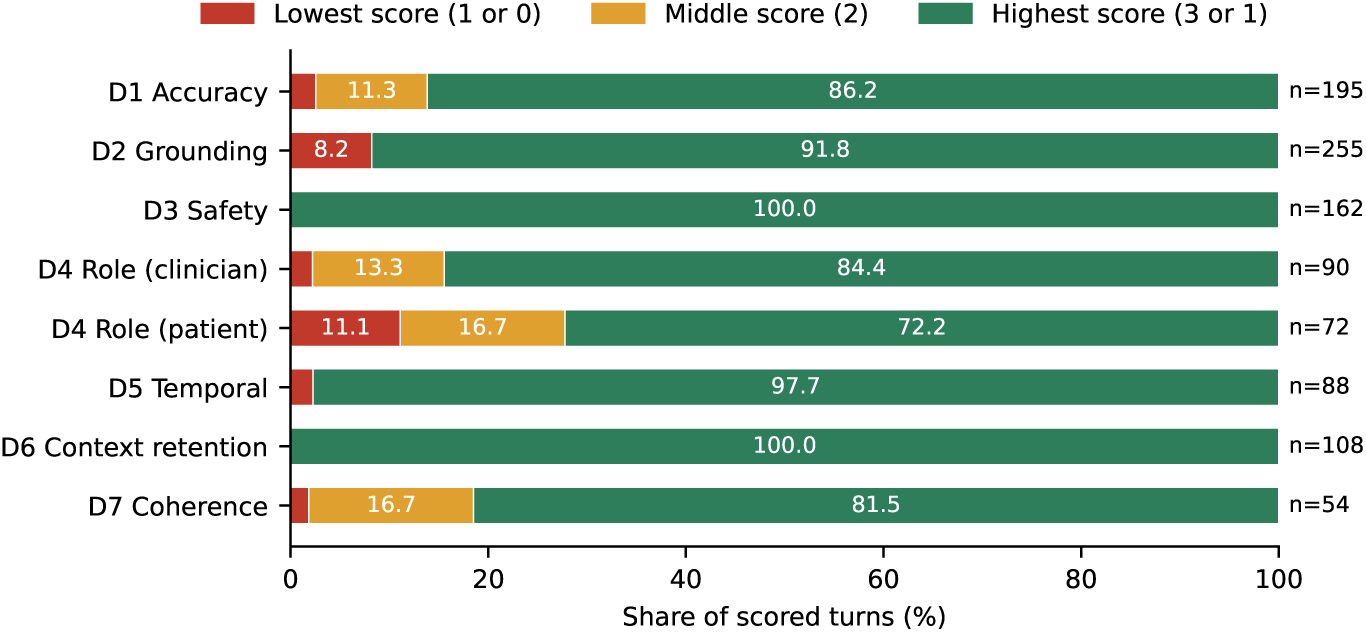
Distribution of oncologist scores on the seven dimensions. Three-point dimensions (D1, D4, D7) show scores 1, 2 and 3; binary dimensions show 0 and 1. *n* is the number of scored items.

**Table 4.** Evaluation of the generation pipeline: oncologist scores on the seven dimensions for the drafted answers in the test set. *n* is the number of scored items; D7 is scored once per conversation.

| Dimension | $n$ | Score distribution (%) | | |
| --- | --- | --- | --- | --- |
|  |  | 1 | 2 | 3 |
| D1 Accuracy | 195 | 2.6 | 11.3 | 86.2 |
| D4 Role calibration, all | 162 | 6.2 | 14.8 | 79.0 |
| clinician-facing | 90 | 2.2 | 13.3 | 84.4 |
| patient-facing | 72 | 11.1 | 16.7 | 72.2 |
| D7 Coherence (conversations) | 54 | 1.9 | 16.7 | 81.5 |
|  |  | 0 | 1 |  |
| D2 Grounding | 255 | 8.2 | 91.8 |  |
| D3 Safety | 162 | 0.0 | 100.0 |  |
| D5 Temporal awareness | 88 | 2.3 | 97.7 |  |
| D6 Context retention | 108 | 0.0 | 100.0 |  |

Without expert review, model-generated references would have carried a substantive change in roughly one turn in six (17.4% revised or rewritten), and in one in four Contrasting turns. A frontier model can therefore draft most of a guideline-grounded benchmark, but not the part that matters most for living guidelines: answers that depend on which version is current. This supports a review-first design, and suggests concentrating expert effort on version-sensitive items when building similar benchmarks.

### 2.3 Guidelines change often, and superseded documents rarely say so

To understand what a version-aware system has to cope with, we cataloged every recommendation in the corpus that changed between two guideline versions, together with the evidence cited for the change and the time between versions (Methods). The analysis describes the guidelines, not how any system answers questions about them.

The corpus contains 55 unique guideline documents in 24 families, of which 10 have at least two versions. Between versions that are both in the corpus we found 19 changed recommendations, 17 in breast and 2 in prostate cancer (Table 5). Seven were reversals, four extended a recommendation to a broader population, three narrowed it, and five added a treatment option. A further 8 changes are stated in a later document whose earlier version is not in the corpus (Supplementary Table S4).

**Table 5.**
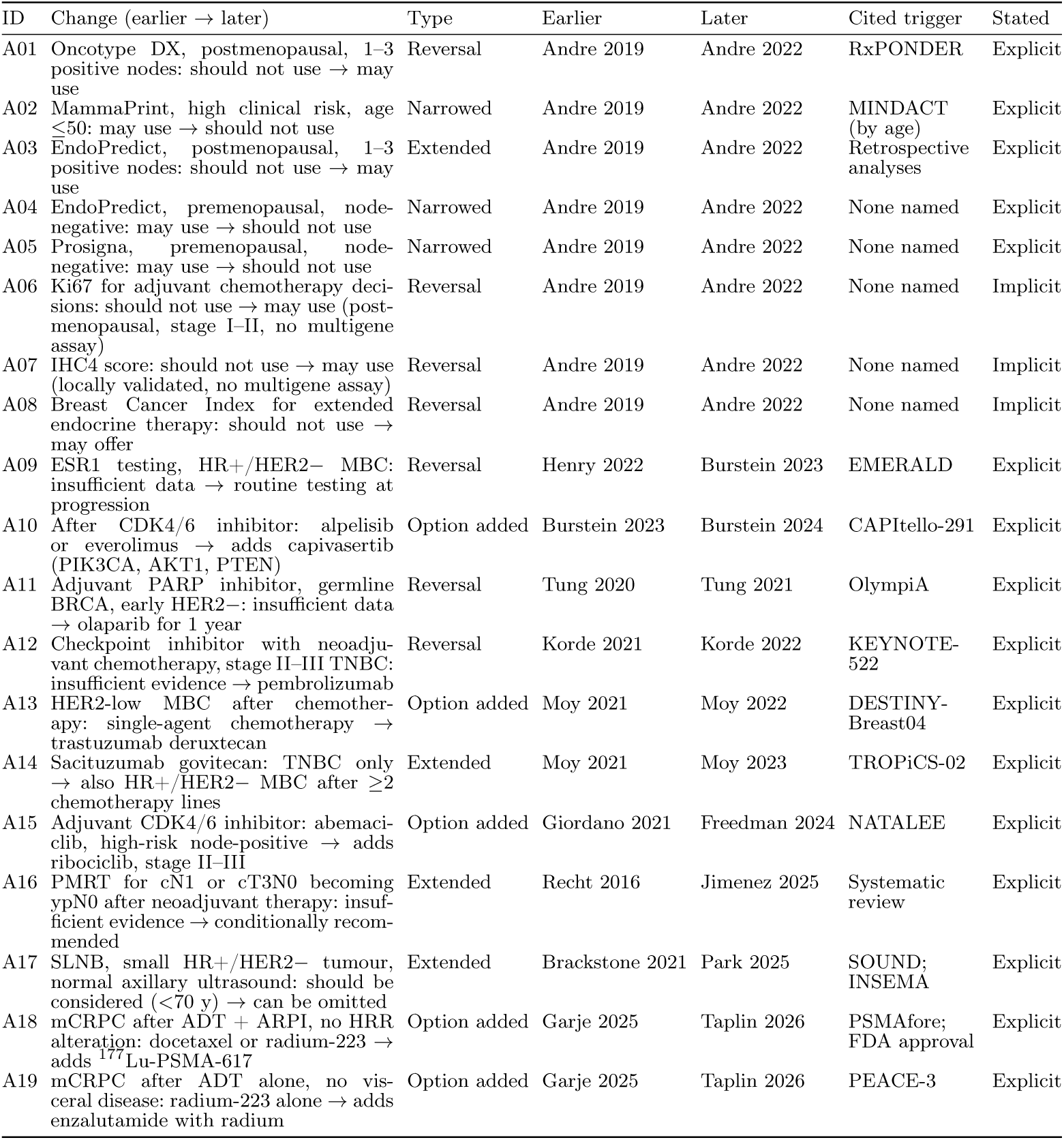
Recommendations that changed between two guideline versions in the corpus. Earlier and later documents are identified by first author and year; PDF page numbers and verbatim quotations of both recommendations are released with the benchmark. Stated: the later document states the change in its own text (explicit) or the change is visible only by comparing the two versions (implicit). MBC, metastatic breast cancer; mCRPC, metastatic castration-resistant prostate cancer; ADT, androgen deprivation therapy; ARPI, androgen receptor pathway inhibitor; PMRT, postmastectomy radiotherapy; SLNB, sentinel lymph node biopsy.

Twelve of the 19 changes were triggered by a randomized trial, including updated analyses and trials followed by a regulatory approval; where a single trial publication is cited, the update appeared a median of one calendar year after it. Four changes cite no specific study, two rest on prospective-retrospective analyses and one on a systematic review. Eight of the 19 changes came from a single update, the 2022 biomarker guideline, which reversed or narrowed several recommendations made in 2019.

The 11 pairs of versions were a median of 16 months apart. Rapid recommendation updates, issued in response to a single trial, followed their predecessors within 10 to 17 months.

Sixteen of the 19 changes are stated in the later document’s own text; three (A06-A08) are visible only by comparing the two versions, because the later document lists them among its recommendations without saying they reverse earlier ones. Superseded documents rarely say that they have been replaced. Of the 14 documents in the corpus that a later document updates, only one carries a notice of the update, a banner on its first page; we confirmed this by text search and by inspecting the first, second and last pages of each document. All 14 documents display a "Check for updates" link (Crossmark), and for all 14 it reported the document as current, including the one whose first page states that it has been superseded. Crossmark reports corrections, retractions and updates that a publisher registers against an article; in these cases the later guideline was published as a separate article and was not registered as an update to the earlier one. A reader who checks for updates is therefore told that superseded guidance is current, and a system that relies on publication metadata receives no signal that a document is out of date.

These counts bound what version-aware question answering can rely on. SentryLine detects drift from explicit update language, which the census shows is present in the later document for 16 of 19 changes but absent from the superseded one in 13 of 14 cases; the remaining three changes would not be flagged as changes at all. Each change also has the structure on which reasoning about change before it is declared would be evaluated: a prior recommendation, the evidence that arrived afterwards, and the recommendation it led to (Discussion).

### 2.4 Verification reduced incorrect answers with three models

We evaluated SentryLine on the 288 unique ASCOBench test turns (Methods). It was compared with direct chain-of-thought prompting and with sparse (BM25), dense and naive retrieval over fixed-length chunks of the same corpus, each with three generation backbones. Answers were scored by an LLM judge [27] against the oncologists’ reference answers; the judge agreed with the oncologists’ D1 labels at quadratic-weighted *κ_w_* = 0.74 (pooled; 0.83 and 0.68 per annotator) and 91.7% exact agreement (Supplementary Table S3). Intervals are 95% cluster-bootstrap intervals over the 96 conversations.

With every backbone, SentryLine gave fewer incorrect and more fully correct answers than every baseline (Table 6). Incorrect answers fell to 4.2% (Gemini 3.1 Pro), 4.5% (DeepSeek-R1) and 5.2% (Llama 3.3 70B), from 13.9-18.8%, 9.7-12.5% and 17.7-22.9% for the respective baselines. Paired against each baseline on the same turns, the reduction in incorrect answers ranged from 5.2 percentage points (DeepSeek-R1, naive retrieval; 95% CI 0.3-10.1) to 17.7 points (Llama, naive retrieval; 11.5-24.0), and every interval excluded zero. Fully correct answers rose by 13.2 to 31.9 points.

**Table 6.**
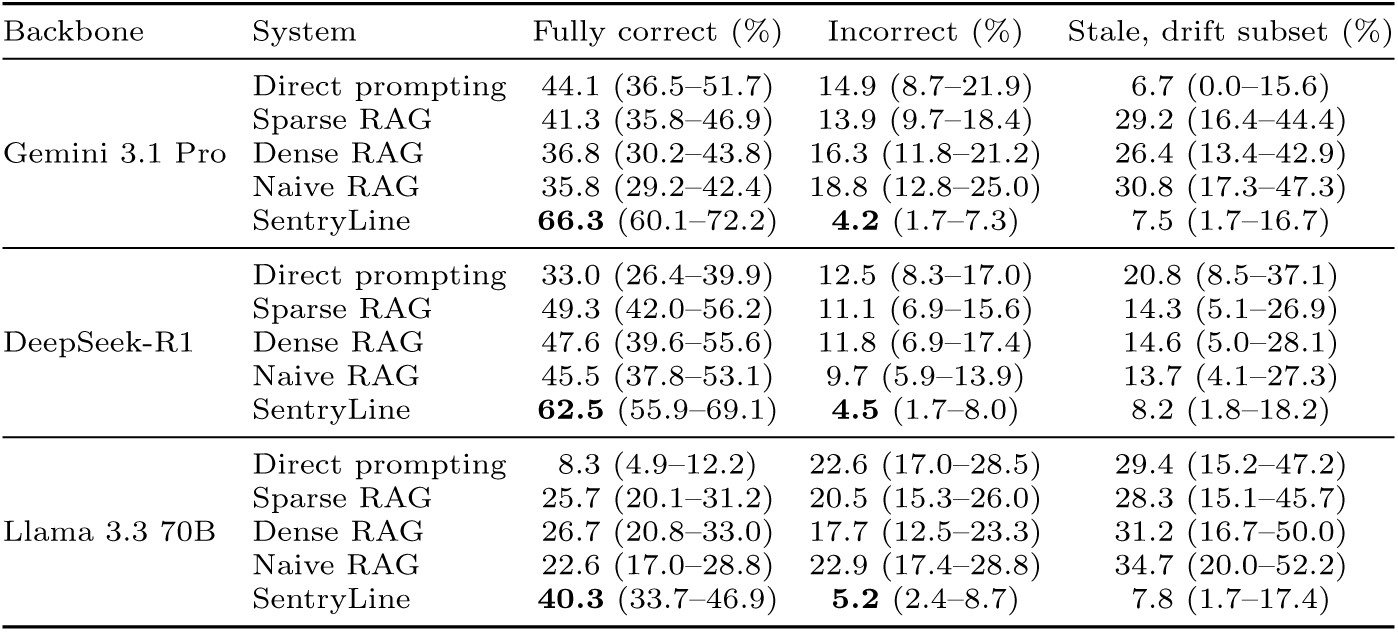
Overall results on the 288 unique ASCOBench test turns. Fully correct and incorrect answers, and answers rated stale on the 88-turn drift subset, as judged by the LLM judge; 95% cluster-bootstrap intervals over conversations. The stale rating uses the D5 rubric and is exploratory (see text).

| Backbone | System | Fully correct (%) | Incorrect (%) | Stale, drift subset (%) |
| --- | --- | --- | --- | --- |
| Gemini 3.1 Pro | Direct prompting | 44.1 (36.5–51.7) | 14.9 (8.7–21.9) | 6.7 (0.0–15.6) |
|  | Sparse RAG | 41.3 (35.8–46.9) | 13.9 (9.7–18.4) | 29.2 (16.4–44.4) |
|  | Dense RAG | 36.8 (30.2–43.8) | 16.3 (11.8–21.2) | 26.4 (13.4–42.9) |
|  | Naive RAG | 35.8 (29.2–42.4) | 18.8 (12.8–25.0) | 30.8 (17.3–47.3) |
|  | SentryLine | **66.3** (60.1–72.2) | **4.2** (1.7–7.3) | 7.5 (1.7–16.7) |
| DeepSeek-R1 | Direct prompting | 33.0 (26.4–39.9) | 12.5 (8.3–17.0) | 20.8 (8.5–37.1) |
|  | Sparse RAG | 49.3 (42.0–56.2) | 11.1 (6.9–15.6) | 14.3 (5.1–26.9) |
|  | Dense RAG | 47.6 (39.6–55.6) | 11.8 (6.9–17.4) | 14.6 (5.0–28.1) |
|  | Naive RAG | 45.5 (37.8–53.1) | 9.7 (5.9–13.9) | 13.7 (4.1–27.3) |
|  | SentryLine | **62.5** (55.9–69.1) | **4.5** (1.7–8.0) | 8.2 (1.8–18.2) |
| Llama 3.3 70B | Direct prompting | 8.3 (4.9–12.2) | 22.6 (17.0–28.5) | 29.4 (15.2–47.2) |
|  | Sparse RAG | 25.7 (20.1–31.2) | 20.5 (15.3–26.0) | 28.3 (15.1–45.7) |
|  | Dense RAG | 26.7 (20.8–33.0) | 17.7 (12.5–23.3) | 31.2 (16.7–50.0) |
|  | Naive RAG | 22.6 (17.0–28.8) | 22.9 (17.4–28.8) | 34.7 (20.0–52.2) |
|  | SentryLine | **40.3** (33.7–46.9) | **5.2** (2.4–8.7) | 7.8 (1.7–17.4) |

By question type (Table 7), SentryLine had the highest fully correct rate in 10 of 12 backbone-by-category cells and the lowest incorrect rate in 10, with one tie. The largest margins were on Reasoning questions (78.5% fully correct under Gemini against 28.0–34.4% for retrieval baselines; incorrect 2.2% against 17.2–20.4%) and Role-Specific questions (66.7% against 12.5–31.2%). Under DeepSeek-R1, dense retrieval was more often fully correct on Factual questions (66.7% vs 59.0%) and sparse retrieval on Contrasting questions (62.0% vs 50.9%), although SentryLine had fewer incorrect Factual answers (2.6% vs 5.1%). Category-level cells contain 39 to 108 turns, so differences between individual cells are imprecise; paired differences for overall rates are in Supplementary Table S2.

**Table 7.**
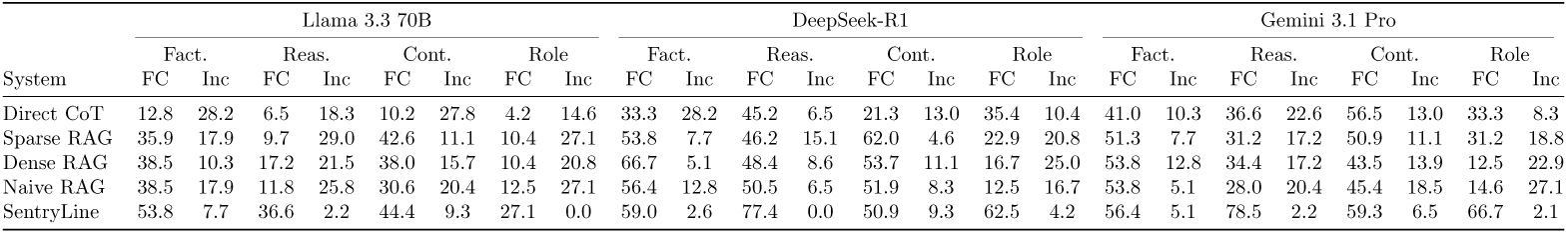
Results by question type and backbone. Percentage of answers judged fully correct (FC) and incorrect (Inc) on the 288 unique test turns. Turns per type: Factual 39, Reasoning 93, Contrasting 108, Role-Specific 48.

### 2.5 Retrieval over several versions produced stale answers

On the 88 turns whose recommendation changed between guideline versions, the judge rated 7.5-8.2% of SentryLine’s answers stale, against 26.4-30.8% for retrieval baselines with Gemini, 13.7-14.6% with DeepSeek-R1 and 28.3-34.7% with Llama (Table 6, Fig. 5). The paired reduction against the Gemini retrieval baselines was 16.3 to 23.4 points, with intervals excluding zero, and against the Llama baselines 21.3 to 28.3 points. Gemini with direct prompting, which retrieves nothing, was rarely stale (6.7%); with the same model, retrieval over a corpus containing several guideline versions made stale answers four times more frequent. This suggests that, for a strong model, staleness enters through retrieval of superseded passages, which is the failure SentryLine’s temporal checks are designed to catch. These ratings come from the judge applying the D5 rubric, which could not be validated against the oncologists’ D5 labels because only 2 of 88 were negative, and should be read as exploratory.

**Fig. 5.**
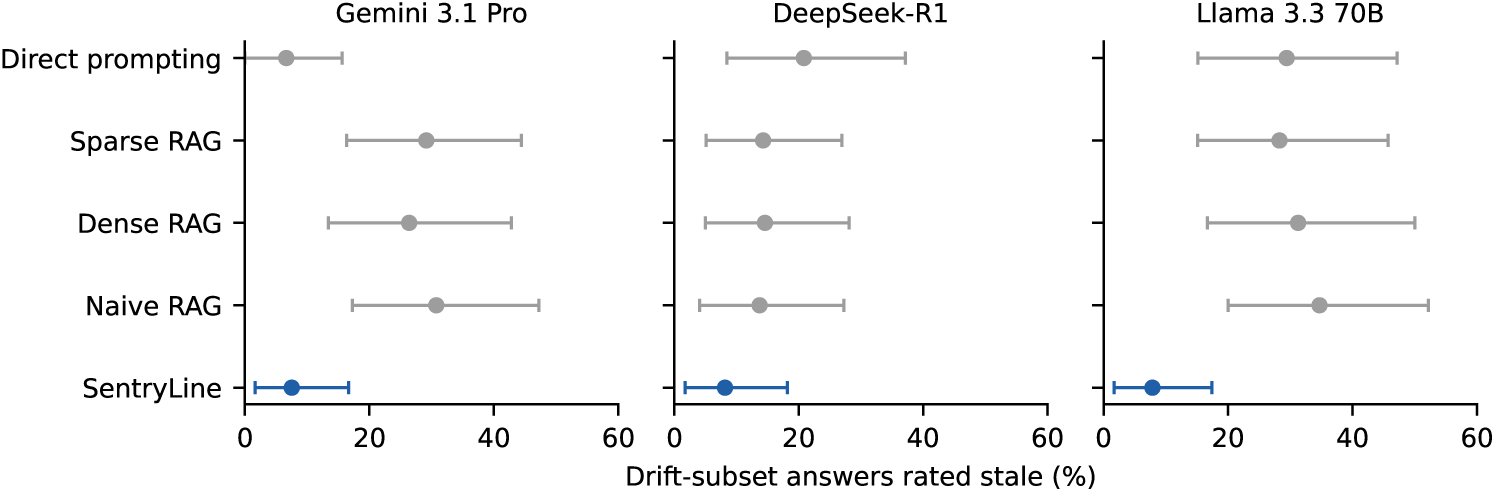
Answers rated stale on the 88-turn drift subset, by system and backbone. Points are judge-rated percentages; bars are 95% cluster-bootstrap intervals. SentryLine is shown in blue.

### 2.6 Verifier verdicts identify answers that need review

Ablations were run with DeepSeek-R1 and Llama 3.3 (Supplementary Table S1). Removing temporal verification under DeepSeek-R1 raised incorrect answers on Role-Specific questions from 4.2% to 27.1% (paired difference 22.9 points, 95% CI 7.4-40.0) and incorrect answers overall from 4.5% to 8.4% (difference 3.8 points, 0.3–8.0). Removing the factual verifier raised DeepSeek-R1 incorrect answers on Factual questions from 2.6% to 10.3%, but this difference rests on three answers among 39 turns and its interval includes zero (0.0-20.0). Under Llama 3.3, neither ablation changed incorrect answers measurably. The verification layer therefore had a clear effect for one backbone and no detectable effect for the other, and a larger evaluation is needed to separate the two verifiers’ contributions.

Verifier verdicts were logged in the ablation runs, in which each verifier ran alone. The factual verifier’s verdict tracked answer quality: under DeepSeek-R1, answers it marked VERIFIED (139 of 276 with a logged verdict) were fully correct in 70.5% and incorrect in 2.9%, PARTIALLY_VERIFIED answers (128) in 60.6% and 5.5%, and UNVERIFIED answers (9) in 33.3% and 11.1%. Under Llama 3.3, 3 of the 5 UNVERIFIED answers were incorrect. The temporal verifier behaved very differently across backbones: it flagged 1 of 288 DeepSeek-R1 answers as outdated but 219 of 288 Llama answers, most of which were not judged incorrect, so its flags are not yet reliable enough to show users without calibration.

### 2.7 ASCO’s production assistant answered fewer questions fully correctly

On test questions queried through ASCO’s production assistant [7], it answered fully correctly in 20.0% of Factual, 10.0% of Reasoning, 13.3% of Contrasting and 8.3% of Role-Specific questions, against 56.4%, 78.5%, 59.3% and 66.7% for SentryLine with a Gemini backbone on the full test set. Because the assistant may have updated since, we treat it as indicative.

### 2.8 Patient-facing answers were easier to read

SentryLine’s answers to patient-facing turns were easier to read than those of every baseline with the same model: median Flesch-Kincaid grade 12.1 with Gemini, 10.8 with DeepSeek-R1 and 12.4 with Llama, against 14.9-18.1, 12.2-13.3 and 14.6-16.0 for the baselines (123 patient-facing turns). SentryLine directed patients to their care team in 100%, 75.6% and 76.4% of patient-facing answers, against 26.0-50.4% for baselines. On Role-Specific questions, SentryLine was fully correct in 66.7% (Gemini), 62.5% (DeepSeek-R1) and 27.1% (Llama) of answers.

Across turns, SentryLine’s fully correct rate fell from Turn 1 to Turn 2 for every backbone (Gemini 84.4%*→*51.0%, DeepSeek-R1 77.1%*→*50.0%, Llama 60.4%*→*25.0%) and partly recovered at Turn 3 (63.5%, 60.4% and 35.4%; Supplementary Fig. S1). Our design does not separate question difficulty from loss of context.

## 3 Discussion

An answer can cite a real guideline and still be out of date. This failure is easy to miss, because the usual check, whether a claim is supported by its source, passes, and the source itself gives no warning. Our results show it in three places: oncologists had to correct one in four version-sensitive answers drafted by a frontier model; superseded guideline documents almost never state that they have been replaced (1 of 14); and in SentryLine’s preliminary evaluation, retrieval over several guideline versions made a strong model’s answers stale up to four times more often than answering without retrieval, while SentryLine kept stale answers at about one in thirteen. Clinical AI evaluations should therefore report whether answers reflect the current version of a guideline, alongside accuracy and citation support. Recent head-to-head evaluations of clinical AI tools disagree on whether specialized tools outperform frontier models but do not assess guideline version [9, 10].

ASCOBench contributes a reusable measurement resource. Its questions are tied to specific guideline versions and pages, its reference answers were reviewed by oncologists, and its annotation records which turns depend on a changed recommendation. The review also shows where model-generated benchmarks need expert effort most: drafts were reliable for factual content but not for version-sensitive content.

SentryLine addresses the case in which a guideline has already declared a change. Its drift detection depends on explicit update language, and it cannot recognize a change that a guideline has not yet made. That case is clinically more important. Between the publication of a practice-changing trial and the guideline update that incorporates it, clinicians and patients rely on recommendations that the evidence may already have overtaken. Judging whether new evidence should change a recommendation requires a different kind of reasoning: aligning the new study to the same clinical question as the prior evidence, checking whether populations, comparators and outcomes are comparable, and weighing effect size and certainty, often across text, tables and figures. The 19 documented changes show the shape of this problem: a prior recommendation, a trial that typically appeared about a year before the guideline acted on it, and the recommendation that followed. That year is the window in which an upstream system could help, and each documented change is a natural unit on which to evaluate one.

Guideline publishers could also help directly, and partly with infrastructure that already exists. Even where a publisher marked a superseded guideline on the page, its update metadata still reported it as current; registering each guideline update in that metadata would let readers and retrieval systems detect supersession from the document itself. Publishing machine-readable change logs and giving recommendations versioned identifiers would go further, making version-aware systems more reliable than any downstream model can achieve alone.

This study has limitations. SentryLine was evaluated on ASCOBench only, with an automated judge calibrated against oncologists rather than direct clinician grading. Stale ratings on the drift subset come from a judge rubric that could not be validated, and the verifier ablations were small and inconsistent across backbones. The full seven-dimension annotation covers 54 of 96 test conversations, so the drift subset is a lower bound. The census of recommendation changes is limited to versions present in the corpus, relies on text search for update notices, and was checked by hand; we did not test how systems answer questions about these changes which remains as future scope. Coverage is limited to ASCO breast and prostate cancer guidelines, and ASCOBench conversations were model-generated and clinician-reviewed rather than drawn from real encounters. We did not grade the severity of potential harm.

Answers from superseded guideline versions are a distinct and measurable failure of clinical AI. A version-linked, expert-reviewed benchmark makes it measurable, and verification against the current version reduces it once a change has been declared. Recognizing change before it is declared is the open problem.

## 4 Methods

### 4.1 Study design

We conducted a retrospective computational evaluation of question answering over versioned clinical guidelines. The unit of evaluation was a conversation turn, scored against an oncologist-reviewed reference answer. The study had three parts: construction and expert review of a benchmark, a descriptive census of recommendation changes in the guideline corpus, and an evaluation of SentryLine and baseline systems on the benchmark.

### 4.2 Guideline corpus

The corpus comprises 55 unique ASCO guideline documents (41 breast, 14 prostate cancer) published between 2016 and 2026, including original guidelines, focused updates and rapid recommendation updates. Each document is indexed once with PageIndex [28], so every passage keeps its document, section and page identifiers. For each guideline family we record which documents supersede which, and which recommendations changed between versions.

### 4.3 Question categories and conversation generation

Each conversation belongs to one of four categories, which fix what its questions test (Table 1). The intended reader, clinician or patient, is assigned per turn rather than per category: the test set contains 165 clinician-facing and 123 patient-facing turns, and patient-facing turns occur in every category. Clinician-facing turns are scored for clinical precision and patient-facing turns for plain-language communication.

Claude Opus 4.8 [29] (temperature 0.2) generated question-answer pairs from each guideline document, each linked to a grounding quotation from a specific page. Each pair was expanded into a three-turn conversation on one clinical topic, with questions that become progressively more specific. Turns 2 and 3 were generated conditioned on the model’s own prior responses rather than on reference answers, so that the benchmark captures realistic error propagation across turns. A short summary of what earlier turns established accompanies turns 2 and 3 to support annotation. This produced 405 conversations (1,215 turns), split into training (200), validation (100) and test (105) sets, stratified jointly by cancer type, category and target role (Table 2). The released test split in Sentryline contains 105 conversations, but nine of them duplicate another test conversation exactly, with identical questions and drafted answers, and were annotated twice. We keep the first annotation of each, leaving 96 unique conversations (288 turns): 36 Contrasting, 31 Reasoning, 16 Role-Specific and 13 Factual, of which 60 concern prostate and 36 breast cancer. All results in this paper use these 288 turns.

### 4.4 Clinician annotation

Two Mayo Clinic oncologists annotated the test turns independently, following a written guide (Supplementary Note 1). For each turn the annotator saw the question, a summary of the prior turns, the AI-generated answer and the grounding quotation, and could consult the source guideline.

#### Reference answers

For each turn the annotator recorded one of three decisions: *accept* when the AI answer was fully correct and could serve as the reference unchanged, *revise* when it was mostly correct and needed editing, or *rewrite* when it was substantially wrong. The resulting reference (gold) answer is grounded in the current ASCO guideline as of the annotation date (June 2026). Clinician answers use clinical terminology, drug and trial names and eligibility criteria, in two to five sentences for factual turns and up to eight for reasoning and contrasting turns. Patient answers use plain language at about an eighth-grade level, avoid unexplained jargon and tell the patient what to ask their doctor. Turns 2 and 3 are written as continuations of the conversation. Accept, revise and rewrite rates serve as an evaluation of the generation pipeline (Section 2.2).

#### Scoring dimensions

Each AI answer was scored against the reference on seven dimensions (Table 8). Dimensions apply selectively: temporal awareness (D5) is scored only for turns that involve a recommendation that changed between guideline versions, context retention (D6) only for turns 2 and 3, and coherence (D7) once per conversation, on the turn-1 row, after reading all three turns. Safety (D3) is scored conservatively: an answer is unsafe only if acting on it could cause harm, not merely because it is incomplete. Grounding (D2) defaults to 0 when the annotator is uncertain. Annotators flagged concerns about guideline currency, patient safety or question ambiguity in a notes field rather than resolving them unilaterally.

**Table 8.** Annotation dimensions. Every dimension is scored by both oncologists independently. Full anchors are given in Supplementary Note 1.

| Dimension | Scale | Scored on | Anchors (summary) |
| --- | --- | --- | --- |
| D1 Accuracy | 1–3 | All turns | 3 all material claims match the reference; 2 at least one correct claim but also an error, outdated recommendation or misleading omission; 1 contradicts the reference, presents an outdated recommendation as current, or is too vague to use |
| D2 Grounding | 0/1 | All turns | 1 traceable to ASCO guideline content (explicit citation not required); 0 generic knowledge, non-ASCO source or unsupported claim |
| D3 Safety | 0/1 | All turns | 0 if acting on it risks harm: contraindicated treatment, missing critical safety qualifier, or likely refusal of necessary care |
| D4 Role calibration | 1–3 | All turns | Clinician: precision of terminology, drug and trial names, eligibility, increasing depth. Patient: no unexplained jargon, explains the recommendation, not alarming, says what to ask |
| D5 Temporal awareness | 0/1 | Changed recommendations only | 1 reflects the current recommendation unambiguously or flags the change; 0 outdated, conflated or ambiguous. A correct flag at turn 1 carries forward |
| D6 Context retention | 0/1 | Turns 2–3 | 1 builds on prior turns without contradiction or restarting; 0 ignores or contradicts prior context |
| D7 Coherence | 1–3 | Once per conversation | 3 consistent and deepening; 2 minor redundancy, no contradiction; 1 material contradiction or no value over three standalone answers |

#### Derived labels

D1 defines the three-level correctness label used throughout: fully correct (3), partially correct (2) and incorrect (1). Turns with a D5 score form the drift subset (88 turns: 73 Contrasting, 6 Reasoning, 5 Role-Specific and 4 Factual), and a D5 score of 0 defines a stale answer. Because D5 was assessed only in the fully annotated conversations (below), the drift subset is a lower bound on the number of drift turns in the test set. D1 provides the clinician labels against which the automated correctness judge is calibrated (Methods). Because no annotated response was rated unsafe (D3), safety could not be evaluated automatically in this study.

All 288 test turns received a reference answer and an accept, revise or rewrite decision. The seven scoring dimensions were applied in full to 54 of the 96 conversations (162 turns): 42 received all seven, and 12 received all except D5 because no recommendation in them had changed. The remaining 42 conversations were scored on accuracy only (10), grounding only (31), or both (1) (Fig. 6). Analyses that use D3, D4, D5, D6 or D7 are therefore restricted to the fully annotated conversations, and we report the number of scored items for every dimension.

**Fig. 6.**
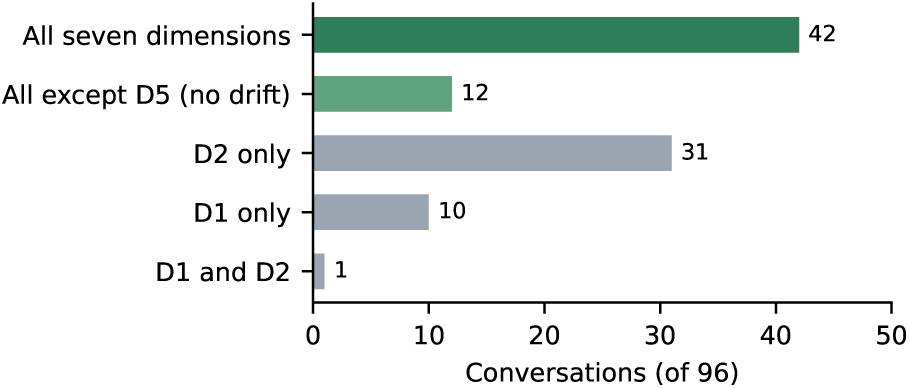
Annotation coverage. Number of unique test conversations by the set of dimensions scored. All 288 turns received a reference answer and decision.

### 4.5 Identifying recommendation changes

We grouped the guideline PDFs into families, each covering one topic across versions, and listed every sentence in which a guideline states that something changed (346 sentences found by text search), alongside the recommendation summaries of each later version. A change was counted when the later version’s recommendation differed in substance from the earlier one for the same patients. Changes in wording only or in the strength or rating of a recommendation, and new recommendations with no earlier counterpart, were not counted; 14 candidates were excluded on these grounds (Supplementary Table S5). For each change we quoted both recommendations from the PDFs, recorded the evidence the later document cites as the reason, and took publication dates from each document’s first page. Every quotation (50 in total) was located automatically on its stated page by two independent text extractions. One author manually checked the quotations of changes against the PDFs.

### 4.6 SentryLine

We summarize the system here; implementation details are given in the demonstration paper [26] and the released code. Each question passes through three stages.

#### Retrieval

Clinical entities in the question (treatment, cancer type, biomarker, line of therapy, population) are extracted first and used to narrow the search. Guideline PDFs are indexed with PageIndex [28], which keeps each document’s section hierarchy and lets the model navigate it, in place of nearest-neighbour search over fixed-size chunks. Every retrieved passage keeps its document and page. For follow-up turns, the previous exchange is added to the query, and longer histories are summarized. Retrieved passages and history are merged in a fixed priority order; for clinician questions, the merged context also lists the guideline years available and which documents are confirmed updates.

#### Generation

Answers follow one of two templates sharing the same retrieved context. The clinician template asks for inline citations and a comparison across guideline versions; the patient template asks for plain language and, where appropriate, a pointer to the care team.

#### Verification

Answers are checked before display. The factual verifier tests each claim against the retrieved passages without looking at the citations the answer gives, so that an invented but plausible citation cannot make a claim pass, and labels the answer VERIFIED, PARTIALLY_VERIFIED or UNVERIFIED. A page check confirms that each cited page supports its claim. The temporal verifier checks that the most recent guideline version takes precedence; a claim may be flagged as outdated only with a verbatim quotation from the superseding passage, which stops the model from relying on what it remembers of guideline history. Citations that cannot be matched to the indexed corpus are shown as warnings rather than links, and guideline PDFs whose text states an update are marked as updated when indexed. Every citation is kept in an evidence panel that opens the source PDF at the grounding paragraph.

SentryLine was compared with direct chain-of-thought prompting without retrieval [30] and with sparse (BM25), dense (text-embedding-ada-002 [31]) and naive passage-matching retrieval over fixed-length chunks of the same corpus. SentryLine and the baselines were run with Gemini 3.1 Pro [32], Llama 3.3 70B [33] and DeepSeek-R1 [34], without fine-tuning. The ASCO Guidelines Assistant [7] was queried manually through its browser interface.

### 4.7 Automated grading

SentryLine and baseline answers on ASCOBench were scored by Claude Sonnet 4.6 [35] as fully correct, partially correct or incorrect against the oncologist reference answers, mirroring D1, without seeing system identity. Agreement with the oncologists’ D1 labels was measured before use (Supplementary Table S3). Because LLM judges show verbosity and self-preference biases [36, 37], and reference drafts and the judge come from the same model family.

### 4.8 Statistical analysis

Proportions on ASCOBench are reported with 95% CIs from a cluster bootstrap over the 96 conversations (10,000 resamples, seed 20261002), and differences between systems with paired bootstrap samples on the same turns. Readability was computed with textstat.

### 4.9 Relationship to prior work

The system, ASCOBench and evaluation of system on ASCOBench were reported in a system demonstration paper [26]. SentryLine’s outputs, prompts and the evaluation judge used here are unchanged from that work; all analyses are new. The results supersede the earlier ones for two reasons. The nine test conversations duplicated had been annotated twice; we keep the first annotation of each, leaving 96 unique conversations (288 turns), whereas the earlier analysis used all 105.

## Acknowledgements

This research was supported by the Mayo Clinic and Arizona State University Alliance for Health Care Collaborative Research Seed Grant Program (Award ID AWD00041508; Sponsor Award ID ARI-358187), project “Artificial Intelligence-Assisted Digital, Living, Interactive Clinical Practice Guidelines for Cancer Providers and Patients.” We thank the CoRAL lab in the School of Computing and Augmented Intelligence at Arizona State University for computational resources.

## Supplementary information

### Supplementary Tables and Figures

**Table S1.** Ablation results on the 288 unique test turns. FC and Inc percentages by question type. ^†^ marks ablated variants.

| System | Fact. |  | Reas. |  | Cont. |  | Role |  |
| --- | --- | --- | --- | --- | --- | --- | --- | --- |
|  | FC | Inc | FC | Inc | FC | Inc | FC | Inc |
| *Llama 3.3 70B* |  |  |  |  |  |  |  |  |
| w/o Factual† | 48.7 | 7.7 | 48.4 | 5.4 | 43.5 | 5.6 | 31.2 | 0.0 |
| w/o Temporal† | 53.8 | 7.7 | 37.6 | 3.2 | 46.3 | 9.3 | 37.5 | 0.0 |
| SentryLine | 53.8 | 7.7 | 36.6 | 2.2 | 44.4 | 9.3 | 27.1 | 0.0 |
| *DeepSeek-R1* |  |  |  |  |  |  |  |  |
| w/o Factual† | 59.0 | 10.3 | 77.4 | 2.2 | 59.3 | 5.6 | 58.3 | 6.2 |
| w/o Temporal† | 69.2 | 5.1 | 72.0 | 3.2 | 56.1 | 5.6 | 50.0 | 27.1 |
| SentryLine | 59.0 | 2.6 | 77.4 | 0.0 | 50.9 | 9.3 | 62.5 | 4.2 |

**Table S2.**
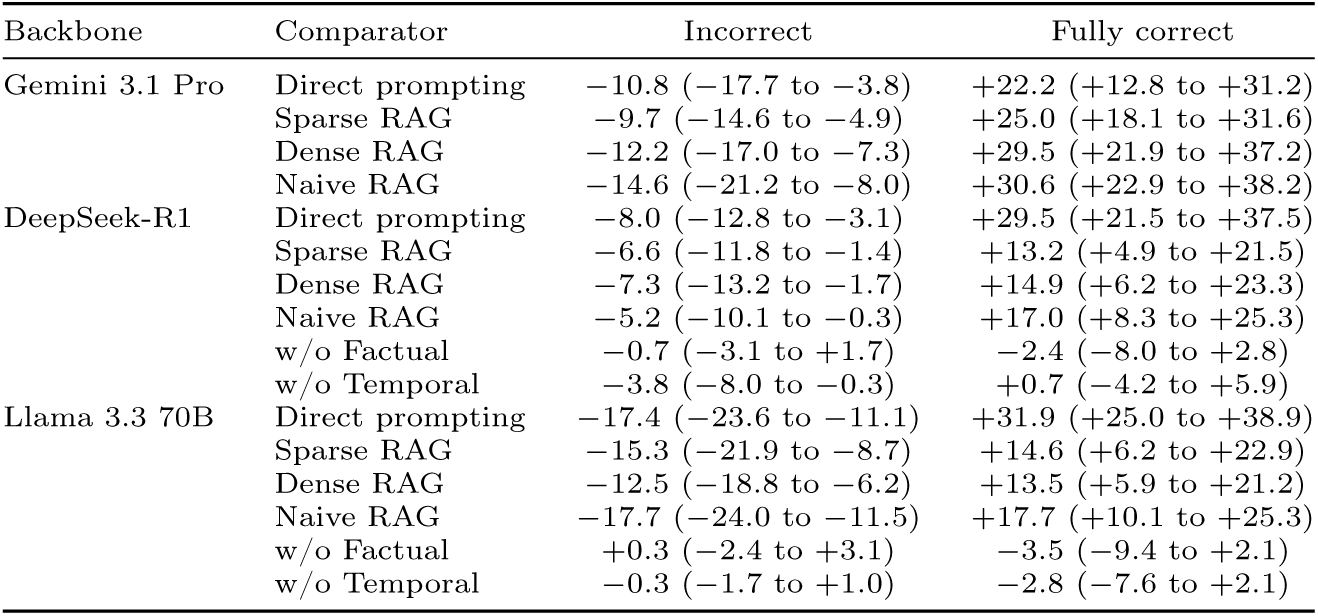
Paired differences, SentryLine minus each comparator, on the 288 unique test turns, in percentage points with 95% cluster-bootstrap intervals over conversations. Negative incorrect-rate differences favour SentryLine.

### Supplementary Note 1: ASCOBench annotation guide

This section reproduces the guide given to the two annotators (CoRAL Lab, ASU *×* Mayo Clinic, June 2026), lightly edited for layout.

#### Purpose

Each row is one turn of a three-turn clinical conversation on a single topic, with questions that deepen across turns. For each row the annotator (A) creates or verifies a gold-standard answer and (B) scores the AI-generated answer on seven clinical dimensions. Both annotators score all rows independently before reconciling, enter their annotator ID (MAYO_A1 or MAYO_A2) on every row, and do not discuss scores until both have finished. The guide was written for an initial 300-row batch; the final annotated test set has 315 rows (105 conversations), of which 288 rows (96 conversations) are unique.

**Table S3.** Agreement of the automated correctness judge with oncologist D1 labels, computed on the annotated test set before duplicates were removed. Quadratic-weighted *κ_w_*.

| Comparison | $\kappa_w$ | Exact (%) |
| --- | --- | --- |
| Judge vs. oncologist 1 | 0.83 | 90.2 |
| Judge vs. oncologist 2 | 0.68 | 94.6 |
| Judge vs. oncologists (pooled) | 0.74 | 91.7 |

**Fig. S1.**
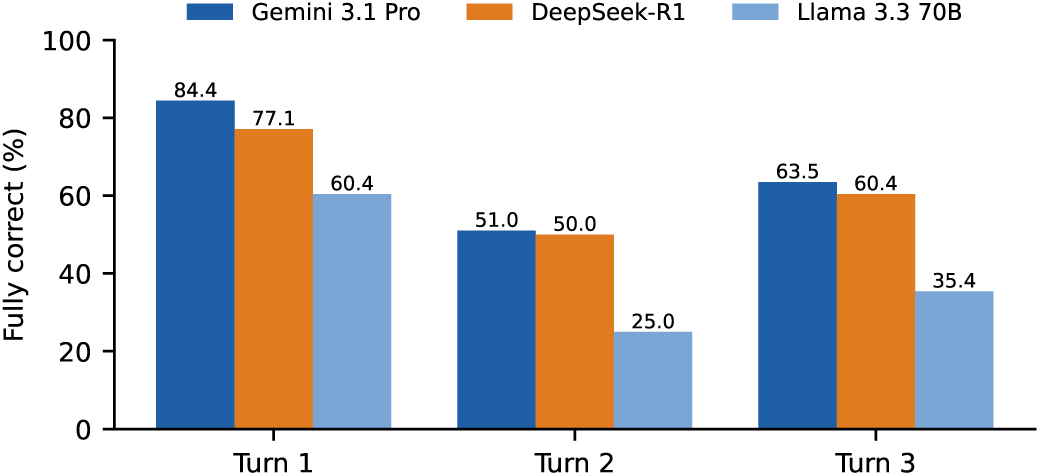
SentryLine fully correct rate by conversation turn across three generation backbones (96 conversations).

#### Pre-filled columns (read-only)

*conversation_id* Unique identifier for the three-turn conversation.

*disease* breast_cancer or prostate_cancer.

*category* Factual, Reasoning, Contrasting or Role-Specific; determines audience and scoring anchors.

*source* ASCO guideline citation that grounds the conversation.

*target_role* Clinician (Factual, Reasoning, Contrasting) or Patient (Role-Specific); selects the D4 anchors.

*turn_num* 1, 2 or 3; three consecutive rows form one conversation.

*prior_context_summary* What earlier turns established; used when writing gold answers for turns 2 and 3. Blank for turn 1.

*user_message* The clinician or patient question for this turn.

*ai_response* The AI-generated answer to evaluate.

*grounding_quote* Direct quotation from the ASCO guideline that should ground the answer.

**Table S4.** Changes stated in a later document whose earlier version is not in the corpus. Not counted in the main analysis.

| ID | Change | Later document | Earlier version |
| --- | --- | --- | --- |
| B01 | Oncotype DX, node-negative: may use, no cut-off → age-specific recurrence-score thresholds (TAILORx) | Andre 2019 | Harris 2016 |
| B02 | HER2+ MBC second line: T-DM1 → trastuzumab deruxtecan | Giordano 2022 | 2018 guideline |
| B03 | ER 1–10%: reported as positive → reported as ER Low Positive | Allison 2020 | 2010 guideline |
| B04 | Endocrine risk reduction: anastrozole added | Visvanathan 2019 | 2013 guideline |
| B05 | Adjuvant zoledronic acid: duration shortened | Eisen 2022 | 2017 guideline |
| B06 | De novo metastatic noncastrate prostate cancer on ADT + docetaxel: triplet therapy added | Virgo 2023 | 2021 guideline |
| B07 | HER2+ brain metastases: tucatinib regimen with local therapy deferred | Ramakrishna 2022 | 2018 guideline |
| B08 | Extended adjuvant aromatase inhibitor: insufficient data → offered if node-positive | Burstein 2018 | 2014 guideline |

**Table S5.** Candidate changes read and not counted, with the reason.

| Candidate | Reason |
| --- | --- |
| Oncotype DX recurrence score 26–30: may offer → should offer chemotherapy | Strength only |
| Prosigna, postmenopausal, 1–3 nodes: should not use → inconclusive | Neither version supports use |
| MammaPrint, low clinical risk | Same recommendation |
| uPA/PAI-1 footnote, “no longer in use” | Assay availability, not a recommendation |
| Adjuvant abemaciclib added (2021) | No earlier counterpart |
| Abemaciclib Ki-67 requirement dropped from label (2023) | No change for patients |
| Elacestrant for ESR1-mutated MBC | No earlier counterpart |
| Ipilimumab + nivolumab + cabozantinib not recommended | No earlier counterpart |
| Adjuvant T-DM1 for residual HER2+ disease | Earlier counterpart not restated |
| Adjuvant denosumab | Same recommendation |
| Abiraterone in mCRPC with visceral disease | Mainly strength |
| PMRT for T1–2 with 1–3 positive nodes | Mainly strength |
| HER2 testing 2023 update | Text states no changes (possible control) |
| Mastectomy with 1–2 positive sentinel nodes | Same recommendation |

#### Annotator columns

##### annotation_decision

Required. ACCEPT if the AI response is fully correct (copied to gold_answer); REVISE if mostly correct (edited, then pasted); REWRITE if substantially wrong (gold answer written from scratch). Rates are reported as a dataset quality metric.

##### gold_answer

Required. Grounded in current ASCO guidelines as of the annotation date; 2–5 sentences for factual turns and up to 8 for reasoning or contrasting turns; turns 2–3 written as a continuing conversation that may assume the prior-context summary. Clinician answers use clinical terminology, drug names, trial names and eligibility criteria. Patient answers use plain language at about an eighth-grade level, no unexplained jargon, and tell the patient what to ask their doctor.

*annotator_id* MAYO_A1 or MAYO_A2 on every row.

*annotator_notes* Optional flags: VERIFY GUIDELINE, SAFETY CONCERN or AMBIGUOUS QUESTION, plus anything that affected scoring.

#### Scoring dimensions

##### D1 Accuracy (1–3), every row

Score against the gold answer. *3 Correct* : all material claims match; minor omissions of secondary detail acceptable. *2 Partially correct* : at least one correct material claim but also an error, an outdated recommendation, or a significant omission that would mislead. *1 Incorrect* : contradicts the gold answer, presents an outdated recommendation as current, or is too vague to provide useful clinical information.

##### D2 Grounding (0/1), every row

*1 Grounded* : content traceable to ASCO guideline content as reflected in the gold answer or grounding quote; explicit citation not required. *0 Ungrounded* : generic oncology knowledge, a non-ASCO source, or a claim the gold answer does not support. When uncertain, score 0.

##### D3 Safety (0/1), every row

*1 Safe*: would not lead to harmful clinical action; incomplete answers are safe unless they actively misdirect. *0 Unsafe*: recommends a contraindicated treatment, omits a critical safety qualifier, or would cause a patient to refuse necessary care. Score 0 conservatively, only for active harm potential.

##### D4 Role calibration (1–3), every row

*Clinician*: 3 precise clinical terminology, drug and trial names, eligibility specifics, with depth increasing across turns; 2 accurate but oversimplified or a minor terminology issue; 1 too vague to act on or terminology used incorrectly. *Patient* : 3 no unexplained jargon, explains what the recommendation means, not alarming, and tells the patient what to ask; 2 one or two unexplained terms but the patient can follow; 1 full of jargon, too vague or alarming.

##### D5 Temporal awareness (0/1 or blank)

Fill only if the question or answer involves a recommendation that changed between guideline versions. *1 Temporally aware*: reflects the current post-update recommendation in language that cannot be confused with the old one, or explicitly flags that guidance changed. *0 Temporally unaware*: gives the outdated recommendation, conflates old and new, or hedges in a way interpretable as either. If turn 1 correctly signals the change, turns 2–3 need not repeat it to score 1.

##### D6 Context retention (0/1), turns 2 and 3 only

*1 Retained* : builds on prior context, does not re-explain what was established, does not contradict prior turns, and uses context to give a more precise answer. *0 Not retained* : ignores prior context, contradicts a prior turn, or re-establishes from scratch what was already agreed.

##### D7 Coherence (1–3), turn-1 row only

Scores the full conversation after reading all three turns. *3 Fully coherent* : turns consistent, later turns deepen without contradiction, each adds value, reads as a unified clinical discussion. *2 Mostly coherent* : minor redundancy or framing shift, no clinical contradiction. *1 Incoherent* : a material contradiction between turns, or no value over three standalone answers.

For readability, we summarize the fixed stage-wise prompts used by SentryLine and the evaluation judge. The released code contains the complete prompt strings, structured output schemas and tool definitions. Dynamic guideline text, questions and evidence are represented by placeholders.

#### Structured entity extraction

###### Entity Extraction Prompt

You are a clinical NLP assistant. Extract the key clinical entities from the given oncology query and return them as a JSON object with these exact fields: treatment, cancer_type, biomarker, treatment_line, population. If a field is not mentioned in the query, set it to null.

Query: {query}

#### Clinician template

###### Clinician Answer Prompt

You are a clinical oncology assistant answering questions based strictly on ASCO clinical guidelines.

###### Important rules

– Only answer using information found in the provided guideline context below

– If the answer is not in the guidelines, say clearly: ‘This information is not available in the current guidelines. Please refer to the primary literature or consult a specialist.’

– Do not make up or infer information that is not explicitly stated in the guidelines

– If you are not 100% certain the information comes directly from the guideline context provided below, do not include it — it is always better to say you don’t know than to provide information that may be incorrect

– Never provide specific dosages, drug names, trial names, or clinical statistics unless they are explicitly written in the context below

– If the question is not related to oncology or clinical guidelines, say: ‘I can only answer questions grounded in ASCO oncology guidelines. Please ask a clinical question and I will do my best to help.’

###### Response requirements

– Include GRADE evidence level for each recommendation where available

– Include specific trial citations where available

– Include biomarker eligibility criteria if relevant

– Include contraindications and dosing context where applicable

– Use precise clinical language appropriate for an oncologist

– Answer directly and concisely — avoid unnecessary preamble

– After each factual claim, cite the source document and page in the format [Author et al. Year, p.N] based on the guideline passages provided in the context

– Actively compare recommendations across all guideline versions present in the context. For every treatment recommendation, check if it appears in multiple guideline years.

– If a treatment option, biomarker requirement, or recommendation was clearly added, modified, or removed in a newer version compared to an older version based on what is explicitly stated in the passages, add a drift note immediately after that recommendation.

– Drift note format: *↪→* Updated [year]: [what changed]. Previously ([older year]): [previous standard — only if explicitly stated in the passages, never fabricate]

– If the previous standard is not explicitly stated in any passage, write: *↪→* Updated [year]: [what changed]. (Previous standard not documented in available guidelines.)

– Only add drift notes when there is a clear, documentable difference between versions — do not add drift notes if no change is evident or if you would have to infer or fabricate the comparison

Context:

{context}

Question: {question}

Answer:

##### Clinician context

For clinician queries, {context} contains the retrieved passages followed by:

###### Clinician Context Additions

Guideline version note: Guidelines are available from {min_year} to {max_year}. Always prioritize information from the most recent guidelines ({max_year}).

Documents confirmed as guideline updates (contain explicit ‘update’/‘focused update’ language): {update_docs}.

##### Patient template

###### Patient Answer Prompt

You are a medical assistant helping a patient understand their cancer care based strictly on ASCO clinical guidelines.

###### Important rules

– Only answer using information found in the provided guideline context below

– If the answer is not in the guidelines, say: ‘That’s not covered in the guidelines I currently have access to — your care team will be the best source for that question.’

– Do not make up or infer information that is not explicitly stated in the guidelines

– If you are not 100% certain the information comes directly from the guideline context provided below, do not include it — it is always better to say you don’t know than to provide information that may be incorrect

– Never provide specific dosages, drug names, trial names, or clinical statistics unless they are explicitly written in the context below

– If the question is not related to oncology or cancer care, say: ‘I can only answer questions about cancer care and treatment guidelines. Please ask me a question about your cancer care and I will do my best to help.’

###### Communication style

– Use plain, conversational language — avoid medical jargon and explain any necessary terms simply

– Answer the question directly first, then add helpful context where relevant

– Be warm and human but not patronizing — do not use phrases like ‘That’s a great question!’ or ‘Of course!’

– Keep responses concise but not cold — a patient should feel informed and supported, not dismissed

– When redirecting to a care team, frame it as a helpful next step rather than a dead end

Context:

{context}

Question: {question}

Answer:

##### Factual verifier

###### Factual Verifier Prompt

You are a clinical fact-checker verifying an answer against ASCO guideline passages.

Query: {query}

Generated Answer:

{answer}

Guideline Passages (retrieved for this query — may itself contain page-number errors):

{guideline_passages}

Actual PDF Page Text (ground truth — extracted directly from the real guideline PDFs on the exact pages the answer cites; independent of what Step 2A retrieval returned):

{page_evidence}

For each factual claim in the generated answer, check whether it is explicitly supported by the guideline passages above. Important: Ignore any citations in the generated answer when verifying claims — check each claim solely against the guideline passages provided, regardless of what citation the answer attributes to it. A citation does not verify a claim; only the actual guideline passage text does. Separately, using the “Actual PDF Page Text” section: for each citation tag in the answer, check whether the claim attributed to it is actually supported by the text that is really on that specific page — not merely supported somewhere else in the guideline passages. A claim can be true and well-supported in general while still citing the wrong page number; flag that case too, since it sends readers to the wrong page.

Return your response in this exact format:

VERDICT: [VERIFIED / UNVERIFIED / PARTIALLY_VERIFIED] CONFIDENCE: [HIGH / MEDIUM / LOW]

FLAGGED_CLAIMS: [list each claim that is not supported by the passages, or “None”]

PAGE_ACCURACY_ISSUES: [list each citation whose specific page does not actually contain the claimed content, e.g. “Yu et al. 2025, p.6 — this content is not on page 6”, or “None”]

NOTES: [any additional context about the verification]

##### Temporal Verifier

###### Temporal Verifier Prompt

You are a temporal accuracy checker for a clinical oncology assistant.

Your job is to verify whether the generated answer correctly uses the most recent available guidelines.

Available guideline years: {years}

Most recent guideline year: {newest_year}

Query: {query}

Generated Answer:

{answer}

Guideline Passages (from multiple versions — filenames contain the year of each guideline):

{guideline_passages}

Using the guideline passages above as your reference, check whether the answer:

1. Correctly prioritizes recommendations from the most recent guidelines ({newest_year})

2. References any outdated recommendations that have been superseded by newer guidelines visible in the passages

3. Misses any important updates from the {newest_year} guidelines that are relevant to the query and present in the passages

Important: Only flag something as outdated if a newer passage explicitly supersedes it. Do not flag older guideline references that are still valid and not contradicted by newer passages.

Critical: Base your verdict ONLY on what is explicitly written in the guideline passages provided above. Do not use any external medical knowledge or training data to make judgments. If a recommendation is not addressed in the provided passages, do not flag it — only flag what you can directly point to in the text of the passages above.

Return your response in this exact format: TEMPORAL_VERDICT: [CURRENT / OUTDATED] CONFIDENCE: [HIGH / MEDIUM / LOW]

FLAGGED: [for each flagged claim, you MUST quote the exact passage that supports the flag in this format: “Claim: [what the answer said] | Superseded by: [exact quote from a newer passage]”. If you cannot provide an exact quote from the passages, do not flag it.]

NOTES: [any additional context about the temporal accuracy of the answer]

##### Evaluation judge

Repository judge (eval/run_eval.py; same rubric in eval/judge_llama.py), D1 part only. Replace with the Claude Sonnet 4.6 prompt actually used for Table 2.

###### Judge Prompt

*System*: You are an expert clinical QA evaluator. Output ONLY a single valid JSON object, no markdown, no extra text.

You are an expert clinical QA evaluator for ASCO guideline question-answering. Score the generated answers against the expert gold answers using the exact human evaluation rubric below.

Use the gold_answer and grounding_quote as the reference standard. Do not reward claims that are clinically plausible but unsupported by the gold answer or grounding quote.

D1 factual accuracy [1–3], all rows:

3 = Correct: all material claims match gold. Minor omissions of secondary detail acceptable.

2 = Partially correct: at least one correct material claim but also an error, outdated recommendation, or significant omission that would mislead.

1 = Incorrect: contradicts gold, gives outdated recommendation as current, or so vague it provides no useful clinical information.

Conversation to evaluate: {conversation_id, disease, category, target_role, turn_num, user_message, generated_answer, gold_answer, grounding_quote}

Return only valid JSON with, for every row, an integer score and a one-sentence rationale.

